# The potyviral protein 6K2 from *Turnip mosaic virus* increases plant resilience to drought

**DOI:** 10.1101/2022.04.04.487062

**Authors:** Ved Prakash, Chad T. Nihranz, Clare L. Casteel

## Abstract

Drought is a major cause of yield loss for crops worldwide. Climate change is predicted to increase global crop losses due to drought through rising temperature and decreased water availability. Virus infection can increase drought tolerance of infected plants compared to non-infected plants; however, the mechanisms mediating virus-induced drought tolerance remain unclear. In this study, we demonstrate *Turnip mosaic virus* (TuMV) infection increases *Arabidopsis thaliana* survival under drought compared to uninfected plants. To determine if specific TuMV proteins mediate drought tolerance, we cloned the coding sequence for each of the major viral proteins and generated transgenic *A. thaliana* that constitutively express each protein. Three TuMV proteins, 6K1, 6K2, and NIa-Pro, enhanced drought tolerance of *A. thaliana* when expressed constitutively in plants compared to controls. Expression of 6K2 also increased plant biomass relative to controls, but had no impact on root biomass, trichome numbers, or on the number of stomata. While drought induced transcripts related to abscisic acid (ABA) biosynthesis and ABA levels in control plants, compared to under well-watered conditions, there were no changes in ABA or related transcripts in plants expressing 6K2 under drought conditions compared to well-watered. 6K2 expression also conveyed drought tolerance in another host plant, *Nicotiana benthamiana*, when expressed using a virus over expression construct derived from *Foxtail mosaic virus* (FoMV). Although the exact mechanisms are still unknown, these results suggest 6K2-induced drought tolerance is ABA-independent and that plant viruses may represent novel sources of drought tolerance for crop plants.

## 1 Introduction

Crops are subjected to biotic and abiotic stresses, which adversely affects their productivity. Drought is one of the most important types of abiotic stress, impacting over three-fourths of global harvested area (approximately 454 million hectares) (Kim, Iizumi & Nishimori 2019). Technological advances have led to improved drought management techniques (Lamaoui, Jemo, Datla & Bekkaoui 2018), however, these technologies often require significant infrastructure. Breeding and transgenic approaches have also lead to the development of drought resistant cultivars, but these plants often have lower yields compared to other cultivars under non-drought conditions (Kim *et al*. 2014; Martignago, Rico-Medina, Blasco-Escámez, Fontanet-Manzaneque & Caño-Delgado 2020). Furthermore, climate change is predicted to intensify the severity and frequency of drought event in the future (Dai 2013), thus identifying new sources of drought tolerance and drought management techniques is a high priority.

While plant viruses are best known as obligate parasites and pathogens of their hosts, recent evidence suggests viruses can enhance host survival during drought. For example, many plants have been shown to perform better under drought conditions when infected with viruses, such *Brome mosaic virus* (BMV), *Cucumber mosaic virus* (CMV), *Cauliflower mosaic virus* (CaMV), *Tobacco mosaic virus* (TMV), *Tomato yellow leaf curl virus* (TYLCV), or *Tobacco rattle virus* (TRV) compared to uninfected control plants (Xu et al. 2008; Westwood et al. 2013; Bergès et al. 2018, 2020; Corrales-Gutierrez et al. 2020; Shteinberg et al. 2021). Virus-induced drought tolerance has been shown using diverse vegetative crops, including rice (*Oryza sativa*), wheat (*Triticum aestivum*), tomato (*Solanum lycopersicum*), pepper (*Capsicum annuum*), watermelon (*Cucumis lanatus*), cucumber (*Cucumis sativus*), and zucchini squash (*Cucurbita pepo*) (Xu et al. 2008; Davis, Bosque-Pérez, Foote, Magney & Eigenbrode 2015). More recently, it has been shown that *Grapevine fanleaf virus* (GFLV)-infected grapevines, a long-lived perennial woody plant, also performs better in mild drought condition compared with the healthy grapevine (Jež-Krebelj et al. 2022). These results suggest viruses may use conserved mechanisms to enhance drought tolerance, and thus understanding the mechanisms of virus-induced drought may enhance our understanding of plant physiology more broadly.

Plants regulate appropriate responses to abiotic and biotic stress using phytohormone signaling pathways. Abscisic acid (ABA) is the primary hormone that regulates plant responses to drought (Wilkinson & Davies 2010), while salicylic acid (SA) is the primary hormone that regulated plant responses to pathogen infections (Ryals *et al*. 1996). CMV-infected *Arabidopsis thaliana* are hypersensitive to ABA, which may contribute to increased drought tolerance (Westwood *et al*. 2013). Aguilar et al. (2017) demonstrated *Plum pox virus* (PPV), expressing the *Potato virus X (PVX)* RNA silencing suppressor molecule P25 (PPV-P25), increases SA levels in host plants. Using transgenic plants expressing *NahG*, they demonstrate PPV-P35 induction of SA is required for virus-enhanced drought tolerance for both *A. thaliana* and *Nicotiana benthamiana* (Aguilar *et al*. 2017). These experiments suggest a role for SA in mediating plant tolerance to drought, in addition to ABA, however the exact mechanisms are still unknown. Recently, it has been shown differences in viral genomes mediate the ability to enhance plant drought tolerance (Westwood et al., 2013; Hily et al., 2016; Carr, 2017; Carr et al., 2018; González et al., 2021). For example, mutating the 2b protein of CMV prevents CMV from conveying drought tolerance and expressing 2b ectopically in host increases drought tolerance of host plants (Westwood *et al*. 2013). These results suggest 2b and other viral proteins could be used as a sources of drought resistance, however to our knowledge, there is only one other drought tolerance conveying viral protein that has been identified so far (Corrales-Gutierrez *et al*. 2020).

Viruses belonging to the genus Potyvirus possess single-stranded positive-sense RNA genomes that are translated into a single polyprotein after host entry and are transmitted by aphids in a non-persistent manner (Revers and García, 2015; Adams et al., 2011; Casteel and Falk, 2016). Recently it was proposed that the function of viral proteins can change in the presence of different ecological interactions and environments (Ray & Casteel 2022). For example, the viral protein Nuclear Inclusion a Protease (NIa-Pro), which is required for the cleavage of the polyprotein into individual mature proteins (Adams, Antoniw & Beaudoin 2005), also has a role in plant-aphid interactions when the aphid vector is present (Bak, Cheung, Yang, Whitham & Casteel 2017). Recent studies have demonstrated that *Turnip mosaic virus* (TuMV) lineages that evolved in the presence of drought increase plant tolerance to drought (González *et al*. 2021). This suggests changes in individual TuMV proteins may convey new functions in drought tolerance under water-limited environments. In this study, we examined additional ecological functions of viral proteins in drought tolerance using TuMV and *Arabidopsis thaliana* as a model. We determined at least three TuMV proteins increase drought tolerance of host plants (NIa-Pro, 6K1, and 6K2), with minimal negative impacts on host biomass. We investigated the underlying mechanisms for one potyviral protein, 6K1, which increased plant biomass under well-watered condition, in addition to increasing drought tolerance in another host species, *Nicotiana benthamiana*. Taken together, our study shows that TuMV 6K2 protein has a new function in host plants under drought conditions, which are independent of the ABA pathway.

## Materials and Methods

### Plants and growth conditions

*A. thaliana* and *N. benthamiana* plants were grown in Lambert LM-111 All Purpose Mix (Quebec, CA) in nursery flats at 25°C and a 16/8 light/dark cycle. The same growth conditions were used in all subsequent experiments. Plants used for all experiments were 4 weeks old, unless otherwise noted. *A. thaliana* plants were used in experiments with TuMV and *N. benthamiana* plants were used in experiments with the FoMV overexpression system.

### Turnip mosaic virus infection of *A. thaliana*

To determine the impact of TuMV on *A. thaliana* drought tolerance, TuMV-GFP was propagated from infectious clone p35TuMVGFP (Lellis, Kasschau, Whitham & Carrington 2002) and used to inoculate 4-week-old *A. thaliana* plants as in Casteel et al., 2015. Briefly, one leaf from each plant was dusted with carborundum and, using a cotton stick applicator, rub-inoculated with sap from a TuMV-GFP-infected *N. benthamiana* plant suspended in 20 mM phosphate buffer. A corresponding set of control plants was dusted with carborundum and mock-inoculated with a cotton stick applicator that was soaked in uninfected sap in 20 mM phosphate buffer. Ten days after inoculation, a UV lamp (Blak Ray model B 100AP, UV Products) was used to identify fully infected leaves. Six-week-old plants were used in drought survival assays unless otherwise noted.

### Stable expression of TuMV proteins in *A. thaliana*

To determine the effect of individual TuMV proteins on *A. thaliana* drought tolerance, we transformed *A. thaliana* individually with one of the ten proteins from TuMV. Individual TuMV proteins (P1, HC-Pro, P3, 6K1, CI, 6K2, VPg, NIa-Pro, NIb, and CP) were previously cloned into the pMDC32 expression vector and recombinant *Agrobacterium tumefaciens* GV3101 generated for each TuMV protein (Curtis & Grossniklaus 2003; Casteel *et al*. 2014). Wild-type *A. thaliana* were transformed with the pMDC32 TuMV protein constructs or the pMDC32 empty vector construct using floral dip transformation (Clough & Bent 1998). Successful transformation was previously confirmed using kanamycin-containing Murashige and Skoog (MS) agar plates (Murashige & Skoog 1962) and then confirmed by reverse-transcription PCR (RT-PCR).

### Ectopic expression of *N. benthamiana* with pFoMV

To further examine the effect of the TuMV protein 6K2 on plant drought tolerance, we cloned the entire coding sequence of *cspB2*, a gene that encodes a bacterial cold shock protein which are known to increase plant tolerance to drought, heat, and salt stress (Castiglioni *et al*. 2008; Nemali *et al*. 2015; Guddimalli *et al*. 2021), and of *cspB2:6K2* into a *Foxtail mosaic virus* (FoMV)-based expression system (Liu *et al*. 2016; Mei & Whitham 2018; Bouton *et al*. 2018). Sequences were cloned into the multiple cloning site I (MCSI) of pFoMV using XbaI and XhoI restriction enzymes. For the cspB2:6K2 construct, a self-cleaving peptide sequence was included to ensure cspB2 and 6K2 are cleaved after translation (Ryan, King & Thomas 1991). Agrobacterium GV3101 was transformed with pCambiaFoMV, pFoMV:cspB2, or pFoMV:cspB2:6K2 and grown individually in Luria-Bertani (LB) with 50μM kanamycin and 10μM rifampicin. The culture was pelleted at 1500 rpm and re-suspended in 10mM of MES buffer containing 10mM MgSO_4_ and 150μM acetosyringone to a final optical density at 600nm (OD_600_). The bacterial suspension was kept in dark for 3 hours and then infiltrated into the underside of the first fully expanded leaf of 4-week-old *N. benthamiana* with a needleless syringe. Infiltrated *N. benthamiana* plants were kept in dark for 24 hours in a growth chamber at 25°C and then transferred to a growth room with a photoperiod of 16/8 h light/dark at 25°C. After one week, infection was verified by RT-PCR.

### Drought stress treatments and survival assays

*Arabidopsis thaliana* drought treatment assays were performed on six-week-old plants. Plants were either subjected to a well-watered or drought treatment. For the well-watered treatment, plants remained in trays saturated with water for 14 days. For the drought treatment, water was withheld from plants for 14 days. After the drought treatment, plants were re-watered and survival was assessed 7 days later. The number of plants that recovered was recorded and the percentage of plants surviving calculated. *Nicotiana benthamiana* drought treatment assays were performed on five-week-old plants. For this experiment, all plants were subjected to a drought treatment. Prior to the start of the drought treatment, potted plants were watered until run-off and then weighted 24 hours later to obtain a baseline mass. Plants were then subjected to a 7-day drought period where soil water content was reduced and maintained at 40% of the baseline pot mass. To maintain the 40% baseline mass, potted plants were weighed each day during the drought period and water was added if the pots fell below the 40% threshold. At the end of the 7-day drought period, plants were re-watered and survival was assessed 4 days later. The number of plants that recovered was recorded and the percentage of plants surviving calculated.

### Water content, biomass, and photosynthetic efficiency measurements

To measure the water content and dry biomass in experiments done with *A. thaliana*, the entire aboveground portion of the plant was cut, fresh weight determined (FW), dried at 37 □ for 7 days, and dry weight determined (DW). Shoot water content was calculated using the following formula: Shoot Water Content = Fresh Weight - Dry Weight / Fresh Weight. To measure root biomass, roots of each plant were carefully removed from the soil by rinsing in water. Roots were then dried at 37°C for one week before weighing. Chlorophyll fluorescence was measured by analyzing the Fv/Fm ratio using a handheld fluorometer (FluorPen FP-110; Photon System Instruments, Brno, Czech Republic). Fv/Fm ratio is commonly used to monitor plant photosynthetic performance and as an indicator of plant stress (Murchie & Lawson 2013; Pérez-Bueno, Pineda & Barón 2019). In experiments with *A*. thaliana, Fv/Fm measurements were taken on the final day of drought and one reading per plant was taken for plants in both the well-watered and drought treatments. In experiments with *N. benthamiana*, Fv/Fm measurements were taken under well-watered conditions (i.e., before drought treatment) and then after 7 days of drought. Three readings per plant were taken during well-watered and drought conditions.

### Stomata and trichome counts

To determine if constitutive expression of 6K2 affected plant morphological traits associated with drought tolerance, we examined the number of leaf trichomes and stomata under well-watered conditions. To quantify the number of trichomes, the adaxial surface of *A. thaliana* leaves were observed using a DinoLite digital microscope (Dunwell Tech, Inc. Torrance, CA) at 55X and the number of trichomes were counted using DinoXcope software (version 2.0.2). To quantify the number of stomata, impressions of the abaxial side of *A. thaliana* leaves were taken using clear nail polish. Leaf impressions were mounted on a slide, observed under a compound microscope at 400X magnification, and total number of stomata was recorded.

### ABA content measurement

To quantify ABA in *A. thaliana* following drought, leaves were collected on the final day of the drought treatment, lyophilized and weighed. Extractions were performed as described previously (Casteel *et al*. 2015). Briefly, leaves were ground into a fine powder and 1 ml of extraction buffer (2:1:0.005 of iso-propanol, HPLC grade H_2_O, and hydrochloric acid) was added to each tube with 1000 ng of D_6_-ABA as an internal control. Samples were shaken in a Harbil paint shaker for 1 min for homogenization. Samples were then centrifuged at 14,000 rpm for 20 min at 4°C. The supernatant was added to a tube containing 1 ml of dichloromethane, vortexed for 30 min at 750 rpm, and centrifuged for 3 min at 12,000 rpm at room temperature. The bottom layer of the solution was transferred to a new tube, dried, and resuspended in 125 ml of methanol. Samples were injected into a Dionex UHPLC system (Thermo Scientific, USA) and ABA detected using a Orbitrap-Q Exactive mass spectrometer (Thermo Scientific, USA). ABA and the internal standard were identified by the signature ions and retention using the Xcalibur 3.0 program (Thermo Fisher Scientific Inc.). Concentrations of ABA were quantified by comparing the peak area of the endogenous compound with the internal standard (deuterated ABA) and expressed relative to weight in mg.

### Total RNA isolation and transcripts abundance analysis

For quantification of ABA- and drought-related gene transcript abundances in *A. thaliana*, leaf samples were collected on the final day of drought treatment, flash frozen in liquid nitrogen, and stored at −80°C until analysis. Each replicate represented a pool of leaf tissue from two individual plants. Total RNA was isolated using Quick-RNA miniprep kit (ZymoResearch, CA, USA) and 1 μg of RNA was used to generate cDNA using SMART MMLV reverse transcriptase (Takara Bio USA, Inc.). Transcript abundance of an ABA biosynthesis gene, *AtNCED3*, and several drought-inducible genes (*AtRD29A, AtDREB19* and *AtRAP2.6L*) were measured using real-time quantitative reverse transcription polymerase chain reaction (qRT-PCR). Expression of *AtNCED3* and *AtRD29* is induced by drought stress (Yamaguchi-Shinozaki, Koizumi, Urao & Shinozaki 1992; Yamaguchi-Shinozaki & Shinozaki 1993; Hikaru *et al*. 2018) while overexpression of *AtDREB19* and *AtRAP2.6L* increases the performance of plants during drought (Krishnaswamy, Verma, Rahman & Kav 2011). qRT-PCR was conducted using SYBR Green PCR Master Mix (Applied Biosystems) and a CFX384™ Optics Module Real-Time System (BioRad Laboratories, Inc.). The qRT-PCR program had an initial denaturation for 2 min at 94°C followed by 40 cycles of 94°C for 30 sec, 55°C for 45 sec and 72°C for 30 sec, and a final extension at 72°C for 5 min. Relative expression was calculated using the delta-delta Ct (2^-ΔΔCt^) method (Livak & Schmittgen 2001). Primers were designed using primer3 for *AtNCED3*-AT3G14440.1 (FP: TCCGGTGGTTTACGACAAGA & RP: TTCCCAAGCGTTCCAGAGAT), *AtRD29A-*AT5G52310.1 (FP: TGAGGAGACGAGAGATGAGAAA & RP: CCGGAGTAACCTAGCATTGAAG), *AtDREB19*-AT2G38340.1 (FP: CTCCTCTGCTTCTGTTGTATCC & RP: CAATCCTTCTTCCTCCCATCTC) and *AtRAP2.6L*-AT5G13330.1 (FP: GAAATCCGCGATCCAAAGAAAG & RP: CAGCTCGGTCATAGGCTAAAG). Ubiquitin was used as internal control (AT4G05320 FP: GGCCTTGTATAATCCCTGATGAATAA & RP: AAAGAGATAACAGGAACGGAAACATA).

### Statistical analysis

Data transformations were performed to meet the assumptions of each statistical test when necessary. Chi-square analysis was used to for survival experiments. Student’s t-tests were performed to assess the effect of 6K2 expression on the quantum efficiency of PSII (i.e., Fv/Fm ratios). One-way ANOVA was used to analyze the effect of 6K2 expression on the number of leaf trichomes, the number of stomata, and the root dry biomass. Two-way ANOVAs were performed to shoot water content, ABA content, and the relative abundance of ABA- and drought-related gene transcripts (i.e., *AtNCED3, AtRAP2.6L, AtDREB19*, and *AtRD29*). The R statistical software (Team 2013) and SAS JMP software (version 16.2.0) were used to perform all statistical analyses.

## Results

### TuMV-enhanced *A. thaliana* survival under drought is mediated by three viral proteins

A greater number of *A. thaliana* plants infected with TuMV survived after 14 days of drought compared to mock-inoculated plants (fig. 1A; 66.21% compared to 25%, respectively; *χ* = 78.0903, *P* < 0.001). To determine if individual TuMV proteins may be mediating enhanced plant survival under drought, survival assays were conducted with *A. thaliana* expressing the 10 major TuMV proteins individually. Expression of 6K1, 6K2, and NIa-Pro increased plant survival after drought treatment compared to plants expressing the empty expression vector (fig. 1B; *χ*^2^ test, *P* < 0.05). Drought resistance can be due to reduced plant size and water needs (Yang *et al*. 2021) and TuMV-infected plants were visually smaller in drought experiments. To determine if this might also be the case for plants expressing 6K1, 6K2, and NIa-Pro, above ground biomass was recorded in a separate experiment without drought. While there was no impact of NIa-Pro expression on shoot biomass compared to controls, unexpectedly, plants expressing 6K1 or 6K2 had a greater biomass than control plants (fig. 1C; one-way ANOVA, TukeyHSD, *F* = 8.15, *P* < 0.05).

**Fig. 1.**
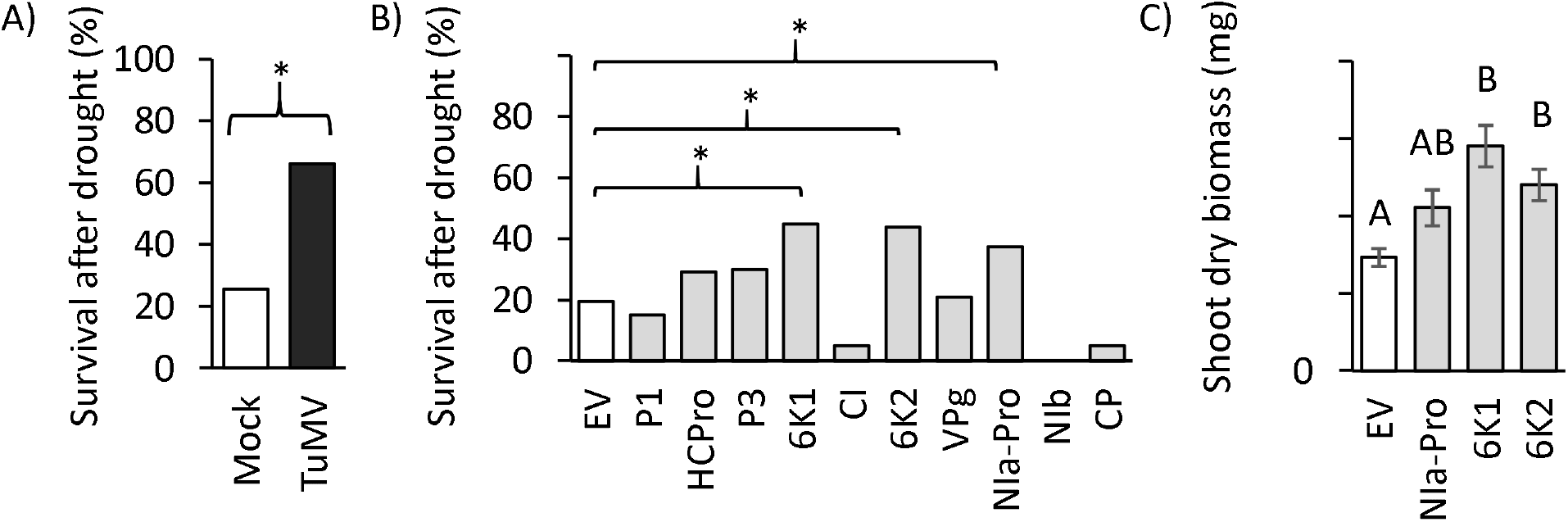
(A) Percent survival of TuMV-infected and mock-infected A. *thaliana* plants following 14 days of drought stress and re-watering (Chi-squire test = 78.0903, *P* < 0.001, N =51-74). (B) Percent of transgenic plants surviving following 14 days of drought stress and rewatering (Chi-squire test, *P* < 0.05, N = 18-87). (C) Dry biomass for 6-week-old *A. thaliana* overexpressing the empty vector (EV), 6K1, 6K2, and NIa-Pro. The entire aerial portion of 6-week-old plants were harvested from the well-watered plants, dried, and weighed (ANOVA, TukeyHSD, *F* = 8.15, *P* < 0.05, N = 10).

### TuMV 6K2 improves the photosynthetic efficiency of *A. thaliana* under drought stress

The 6K2 protein localizes to the chloroplasts of plant cells (Taiyun *et al*. 2010; Cheng *et al*. 2021) and several chloroplast proteins have be linked to increased drought resistance (Lim, LEE & JANG 2014; Nawaz, Lee, Park, Kim & Kang 2018); therefore, we decided to investigate plants expressing 6K2 in more detail. First, we measured chlorophyll fluorescence (i.e., Fv/Fm ratios) in 6K2 overexpressing plants and controls (empty-vector transformed) under well-watered and drought conditions. Fv/Fm ratios were not significantly different between empty vector control plants and 6K2 plants under well-watered condition, however, during drought stress, the photosynthetic efficiency of 6K2 transgenic plants was significantly higher than empty vector control plants (fig. 2A, *t*-test; *t* = 7.6199, df = 7.6199, *P* = 0.017). We also measured shoot water content of EV control and 6K2 overexpressing plants under well-watered and drought conditions. 6K2 overexpressing plants had greater shoot water content than EV control plants in well-watered condition (fig. 2B; two-way ANOVA, TukeyHSD, *F* =17.13, *P* < 0.05). However, during drought, there was no significant difference in the shoot water content between 6K2 overexpressing plants and EV control plants (fig. 2B; two-way ANOVA, TukeyHSD, *F* =17.13, *P* > 0.05).

**Fig. 2.**
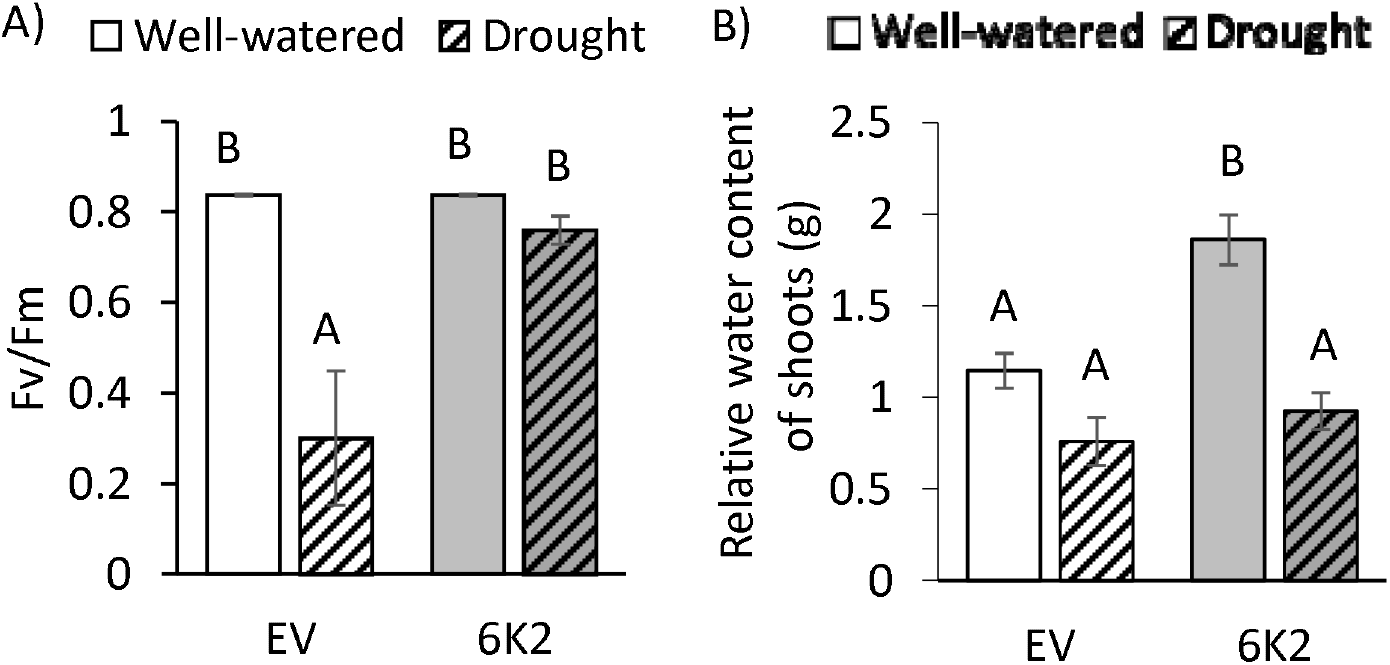
(A) Fv/Fm ratio of the dark-adapted leaves was measured for transgenic *A. thaliana* plants expressing the TuMV protein 6K2 or the empty expression vector (EV) under well-watered conditions and after one week of drought stress (*t*-test, *P* < 0.05, N=8). (B) Relative shoot water content was calculated fresh weight at the end of the experiment using the following formula: (fresh weight (FW) – dry weight (DW))/ FW. (Two-way ANOVA, TukeyHSD, *F* =17.13, *P* < 0.05, N = 8). Letters represent significant difference.

### The 6K2 protein does not alter physical drought tolerance traits in *A. thaliana* plants

Several physical plant traits have been associated with enhanced drought tolerance, such as increase cuticular wax (Guo *et al*. 2016), root biomass (Klein, Schneider, Perkins, Brown & Lynch 2020), trichome numbers (Bosu & Wagner 2007), and stomata (Lecoeur, Wery, Turc & Tardieu 1995; Galmés, Flexas, Savé & Medrano 2007). To determine if 6K2 expression may be increasing drought tolerance through physical mechanisms, we next measured plant root biomass and the number of leaf trichomes and stomata. There was no significant difference in root biomass (fig. 3A; one-way ANOVA, *F* = 0.282, *P* > 0.05), the number of trichomes (fig. 3B; one-way ANOVA, *F* = 1.284, *P* = 0.269), or number of stomata (fig. 3C; one-way ANOVA, *F* = 1.035, *P* = 0.317) between plants expressing 6K2 compared with control plants.

**Fig. 3.**
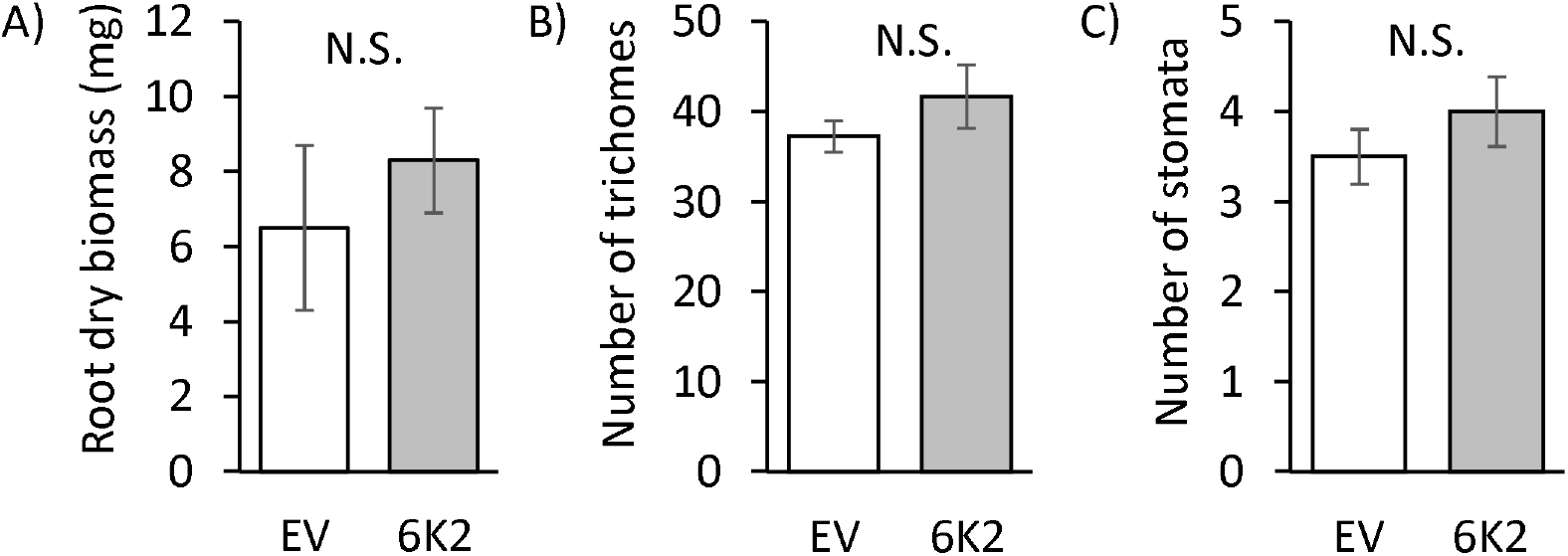
(A) Root dry biomass of well-watered A. *thaliana* overexpressing the TuMV protein 6K2 or the empty expression vector (EV) (one-way ANOVA, *F =* 0.282 *P* = 0.6026, N=10). (B) Number of trichomes measured in a single field at 55X (one-way ANOVA, F = 1.284; *P* = 0.269, N=12) and (C) stomata measured in a single field at 400X (one-way ANOVA, F = 1.035, *P*=0.317 N=16) in well-watered 6K2 overexpressing lines of *A. thaliana* and EV *A. thaliana*.

### TuMV 6K2 protein provides drought tolerance to *A. thaliana* independent of ABA pathway

Plants increase ABA in response to drought stress, which leads to physiological changes, such as regulating water status, activation of genes required for dehydration resistance, and elevating plant antioxidant defense system (Zhou *et al*. 2019). Thus, we hypothesized that ABA content may be elevated in plants expressing the 6K2 protein relative to controls. To address this we measured the abundance of *AtNCED3*, a transcript related to ABA biosynthesis (Bhaskara, Nguyen & Verslues 2012; Hikaru *et al*. 2018) and ABA levels in plants under well-watered and drought conditions. *AtNCED3* and ABA content drastically increased in the drought-treated EV control plants compared with EV plants under well-watered condition, which was expected (fig. 4A, two-way ANOVA, TukeyHSD, *F* = 17.2120, *P* < 0.05; fig.4B; two-way ANOVA, TukeyHSD, *F* = 13.4817, *P* < 0.05). However, there was no significant change in *AtNCED3* transcripts or ABA content of the 6K2 plants in well-watered or drought conditions compared to controls (fig. 4A-B). Taken together, these results suggest that TuMV 6K2 transgenic plants are either insensitive to drought or increasing drought tolerance in ways that are independent to ABA signaling.

**Fig. 4.**
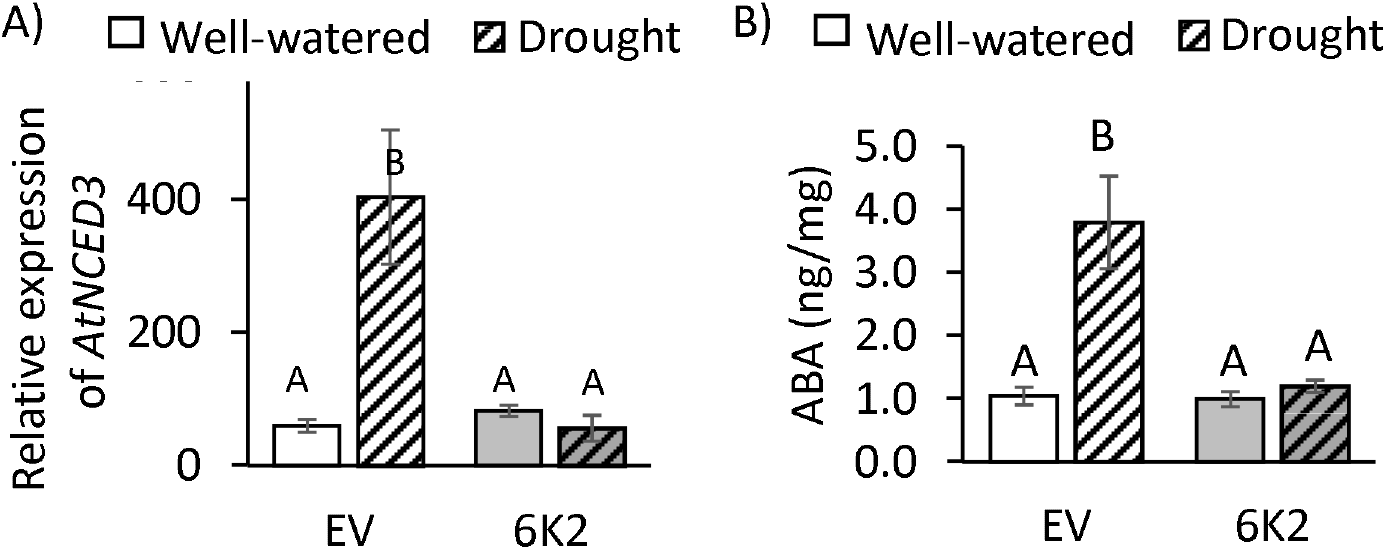
(A) Relative abundance of *AtNCED3*, a transcript related to ABA biosynthesis, and (B) ABA levels in *A. thaliana* plants expressing the empty vector (EV) or the TuMV protein 6K2 under well-watered and drought conditions. *AtUbiquitin* was used as an internal control and each replicate was a pool of leaves from two individual plants. For transcript abundance: N = 5, two-way ANOVA, TukeyHSD, *F* = 17.2120, *P* <0.05). For ABA levels: N = 5, two-way ANOVA, TukeyHSD, *F* = 13.4817, *P* <0.05). Letters indicate significant differences.

### *A. thaliana* plants overexpressing TuMV 6K2 do not induce drought-associated transcripts

Previous studies show that the expression of *AtRD29* is induced by drought and ABA (Yamaguchi-Shinozaki *et al*. 1992; Yamaguchi-Shinozaki & Shinozaki 1993), and overexpression of *AtRAP2.6L* and *AtDREB19* enhances the performance of the plants under drought (Krishnaswamy *et al*. 2011). Although all three transcripts increased in EV plants under drought conditions compared to well-watered conditions, there were only significant differences for *AtRD29* (fig. 5A-C; two-way ANOVA, TukeyHSD, for *AtRD29, F* = 8.0594, *P* < 0.05; *AtRAP2.6L, F* = 0.3477, *P* = 0.77; for *AtDREB19, F* = 3.0379, *P* = 0.31). Similar to *AtNCED3* and ABA, we did not observe significant difference in the accumulation of any of the three transcripts in well-watered or drought condition for plants expressing 6K2 compared to controls (fig. 5A-C).

**Fig. 5.**
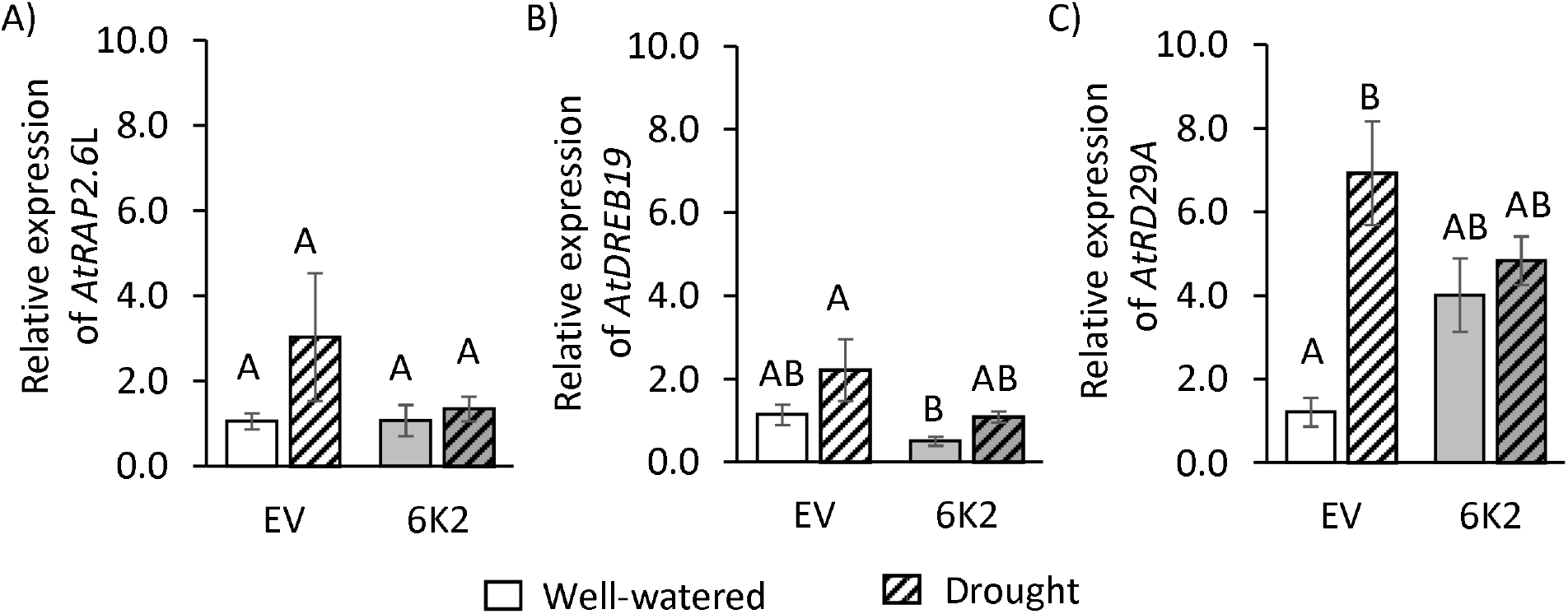
Relative abundance of three drought responsive genes (A) *AtRAP2.6L*, (B) *AtDREB19*, and (C) *AtRD29A* in *A. thaliana* expressing the empty vector (EV) or the TuMV protein 6K2 under well-watered and drought conditions. *AtUbiquitin* was used as an internal control and each replicate had leaf samples pooled from two individual plants. For transcript abundance: N = 5, two-way ANOVA, TukeyHSD (for *AtRAP2.6L, F* = 0.3477, *P* >0.05; for *AtDREB19, F* = 3.0379, *P*>0.05; for *AtRD29, F* = 8.0594, *P* <0.05). Letters indicate significant differences.

### TuMV 6K2 provides drought tolerance to *N. benthamiana*

A pFoMV vector was constructed previously to express *cspB2*, a bacterial cold shock protein known to increase plant tolerance to drought stress (Castiglioni *et al*. 2008; Nemali *et al*. 2015; Guddimalli *et al*. 2021). To evaluate if 6K2 expression enhances *cspB2* drought tolerance and also can be used in other host plants, we modified this pFoMV:cspB2 construct to also express 6K2 (i.e., pFoMV: cspB2:6K2) and assessed survival and photosynthetic efficiency of plants infected with the constructs following a drought treatment. We found that *N. benthamiana* plants infected with pFoMV:cspB2:6K2 had greater survival than plants infected with pFoMV (fig. 6A; *χ* = 0.681, *P* = 0.056) and pFoMV:cspB2 (fig. 6A; *χ* = 4.557, *P* = 0.033). However, there was no difference in survival between plants infected with pFoMV and pFoMV:cspB2 (fig. 6A; *χ* = 0.085, *P* = 0.771). Furthermore, following a 7-day drought treatment, the photosynthetic efficiency of *N. benthamiana* infected with pFoMV:cspB2:6K2 was greater than plants infected with pFoMV (fig. 6B; *t* = −2.627; df = 13.908, *P* = 0.02) and pFoMV:cspB2 (fig. 6B; *t* = − 2.4504, df = 10.78, *P* = 0.033). No significant differences in photosynthetic efficiency were observed between plants infected with pFoMV and pFoMV:cspB2 under drought conditions (*t* = 0.5933; df = 11.317, *P* = 0.565) or among any treatments under well-watered conditions (fig. 6B).

**Fig. 6.**
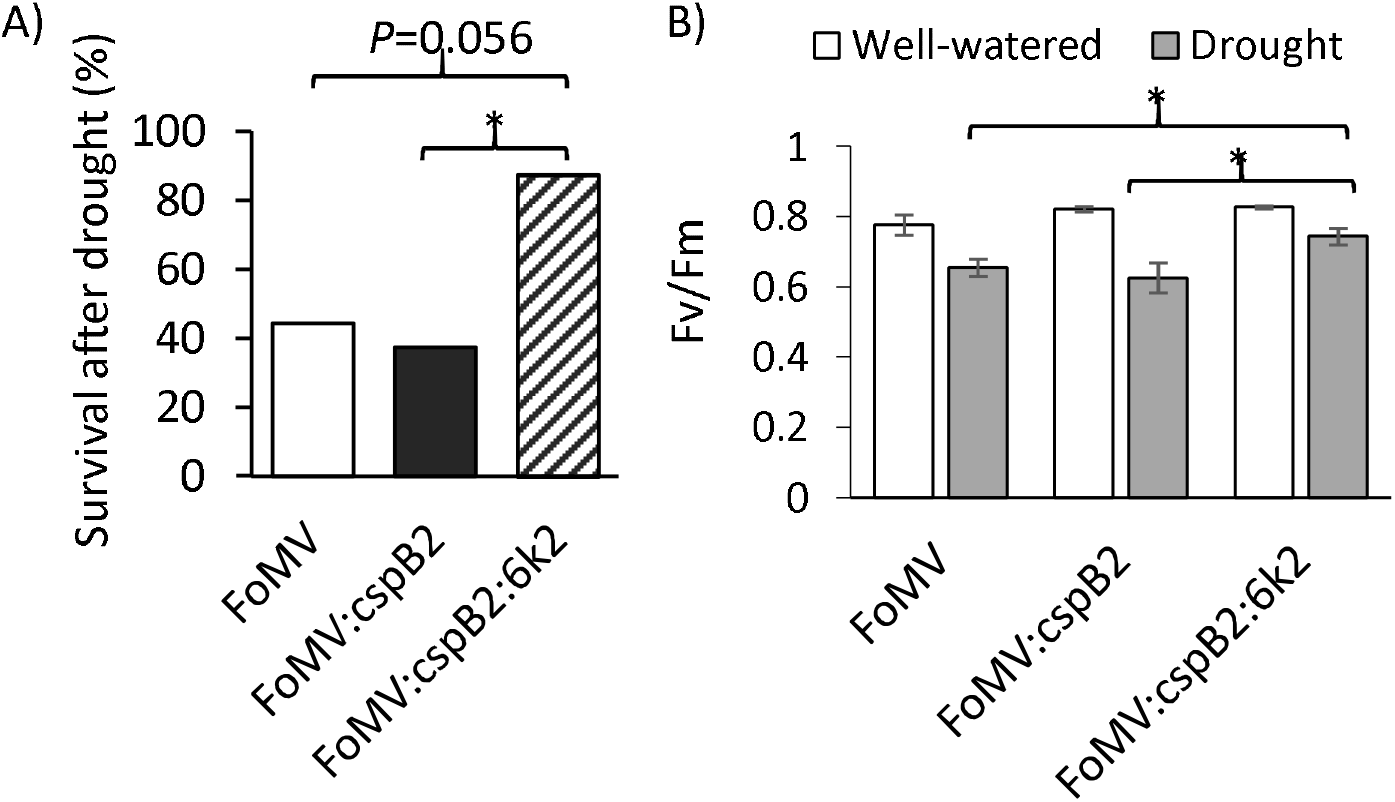
(A) *N. benthamiana* plants were infected with pFoMV, pFoMV:cspB2, or pFoMV:cspB2:6K2 and percent survival recorded following 7 days of drought stress and rewatering (Chi-squire test, * indicates *P* < 0.05, N = 8). (B) Fv/Fm ratio of the dark-adapted leaves was measured from plants after one week of drought stress (Student’s t-test, * indicates *P*<0.05, N=8).

## Discussion

In this study, we demonstrate TuMV can benefit its host *Arabidopsis thaliana* by increasing plant survival under drought conditions (fig. 1A). We go further and show at least three TuMV proteins, 6K1, 6K2, and NIa-Pro, are responsible for enhanced drought tolerance in TuMV-infected plants (fig. 1B), although the exact mechanism of how each potyviral protein enhances drought resistance is still unknown. While bacterial cold shock proteins have been shown to increase plant drought tolerance (Castiglioni *et al*. 2008; Yu *et al*. 2017), we found that expression cspB2 alone in *N. benthamiana* did not increase survival or alleviate drought induced changes to the photosynthetic efficiency of *N. benthamiana* plants. Expression of cspB2 with TuMV 6K2, however, increased both *N. benthamiana* survival and photosynthetic efficiency following a drought treatment compared to controls (fig. 6A,B). Due to the small size of most viral genome, many viral proteins have evolved multifunctionality (Callaway, Giesman-Cookmeyer, Gillock, Sit & Lommel 2001; Deng, Wu & Wang 2015; Valli, Gallo, Rodamilans, López-Moya & García 2018). The ability of 6K1, 6K2, and NIa-Pro to enhance drought tolerance may be related to the proteins primary function, or alternatively, it may be an additional ecological function (Ray and Casteel, 2022). Recently it was determined that the potyviral NIa-Pro has evolved the ability to inhibit plant defenses only when insect vectors are present (Bak *et al*. 2017), which should benefit the virus through enhanced transmission (Ray and Casteel, 2022). It is tempting to speculate that drought tolerance-inducing viral proteins will benefit the virus as well as the host, as greater host survival under drought conditions would increase the time viruses can replicate and spread in the environment before the host dies.

Virus infection causes molecular, physiological and biochemical changes in plants that can induce a state of drought tolerance (Xu *et al*. 2008; Hily *et al*. 2016; Carr *et al*. 2018; González *et al*. 2021). In our study, expression of 6K2 increased plant biomass relative to controls (fig. 1C), yet still prevented drought induced changes in photosynthetic efficiency (fig. 2A). There was no impact of 6K2 expression on root biomass, trichome numbers, or on the number of stomata of *A. thaliana* (fig. 3A-C). ABA regulates stomatal conductance and expression of various genes, such as *AtRD29*, *AtDREB19* and *AtRAP2.6L* which improves plant performance under drought (Iuchi *et al*. 2001; Krishnaswamy *et al*. 2011; Jia, Zhang, Ruan, Wang & Wang 2012; Munemasa *et al*. 2015). To our surprise, there was no change in ABA biosynthesis transcripts (*AtNCED3*) or ABA content in the 6K2 overexpressing *A. thaliana* under drought compared with the non-drought condition, while in controls plants ABA biosynthesis transcripts and ABA content drastically increased under drought (fig. 4). Although expression of *AtRD29*, *AtRAP2.6L* and *AtDREB19* increased in control plants under drought, this was only significant for *AtRD29* (fig.5) and none of these transcripts increased in *6K2* overexpressing *A. thaliana* leaves under drought compared to well-watered conditions (fig. 5). Taken together, these results suggests that TuMV 6K2 transgenic plants are either insensitive to drought, or drought tolerance was increased in ways that are independent of ABA signaling.

A similar study has shown that C4 protein of *Tomato yellow leaf curl virus* (TYLCV) provides drought tolerance in *A. thaliana* via an ABA-independent pathway (Corrales-Gutierrez *et al*. 2020). It has recently been suggested that the ability of TuMV to enhance drought tolerance in the Oy-1 accessions of *A. thaliana* involves the downregulation of ABA-independent gene expression pathways (González *et al*. 2021), further supporting our findings. Although the same differential gene expression patterns were not observed for other TuMV-infected *A. thaliana* accessions under drought, suggesting the mechanisms used by TuMV to enhance drought tolerance may be dependent on plant genotype (González *et al*. 2021). In our study, 6K2 expression enhanced survival of two different plant species under drought conditions (*A. thaliana*: fig. 1B; *N. benthamiana*: fig. 6A). These experiments suggest the mechanisms used by 6K2 to enhance drought tolerance may be somewhat conserved. Furthermore, the role of subclass I SNF1-RELATED PROTEIN KINASES 2 (SnRK2s) has been linked with ABA-independent drought stress tolerance in plants (Soma, Takahashi, Yamaguchi-Shinozaki & Shinozaki 2021). Thus, it would be interesting to evaluate the activity of these protein kinases in virus infected plants to further understand the mechanism of drought tolerance and if 6K2 affects its expression or activity.

TuMV has also been reported to decrease drought tolerance of host plants (Manacorda *et al*. 2021). In this work, water was withheld from *A. thaliana* plants immediately after the plants were infected with TuMV, thus infection was not established in host plants before drought. Manacorda et al. (2021) found that plants that were infected with TuMV immediately before drought had higher mortality compared to control plants. We used the same plant and virus genotype in our study, however we allowed virus infection to establish for two weeks before exposing plants to drought. The inability of TuMV to increase host drought tolerance at early stages in the infection process may be due to 6K2 and 6K1 playing crucial roles in viral replication and infection establishment (Taiyun & Aiming 2008; Laliberté & Sanfaçon 2010; Cui & Wang 2016). NIaPro is required for the cleavage of the viral polyprotein into individual mature proteins, another essential function in establishing infections (Adams *et al*. 2005). These primary functions of 6K1, 6K2, and NIa-Pro may have to be prioritized over enhancing drought tolerance to ensure a successful infection, although additional experiments would be needed to test this.

In the field, crops often experience a combination of abiotic stresses which cause reduced yield and enormous economic loss. While our study demonstrates that TuMV and specific TuMV proteins increase plant survival under drought, other aspects of plant-virus interactions and environmental conditions may influence these impacts. For example, when Arabidopsis was subjected to drought and heat stress, plants became more susceptible to TuMV infection (Prasch & Sonnewald 2013). Under drought condition, TuMV accumulation may be limited to a certain threshold to ensure host survival. However, the combined impact of heat and drought stress may break this threshold with unknown impact on host drought tolerance. Clearly, additional work is still needed, and our study and others help set the stage for advancing our understanding of the plant-microbe interactions in complex environments.

## Author Contributions

CLC conceived the project. VP, CTN, and CLC designed the research. VP, CTN, and CLC performed the research, analyzed the data and interpreted the data. CLC and VP wrote the article with contributions from CTN.

## Acknowledgments

We thank Dr Swayamjit Ray for helping in LC-MS/MS. We thank Dr Georg Jander for his help in lyophilization. This work was supported by Defense Advanced Research Projects Agency (DARPA) agreement HR0011-17-2-0053 and US National Science Foundation award #1723926 to C.L.C.

## References

Adams M.J., Antoniw J.F. & Beaudoin F. (2005) Overview and analysis of the polyprotein cleavage sites in the family Potyviridae. Molecular plant pathology 6, 471–487.

Adams M.J., Zerbini F.M., French R., Rabenstein F., Stenger D.C. & Valkonen J.P.T. (2011) Family potyviridae. Virus Taxonomy, Ninth Report of the International Committee on Taxonomy of Viruses (AMQ King, MJ Adams, EB Carstens & EJ Lefkowitz, eds), 1069–1089.

Aguilar E., Cutrona C., del Toro F.J., Vallarino J.G., Osorio S., Pérez-Bueno M.L., … Tenllado F. (2017) Virulence determines beneficial trade-offs in the response of virus-infected plants to drought via induction of salicylic acid. Plant, Cell & Environment 40, 2909–2930.

Bak A., Cheung A.L., Yang C., Whitham S.A. & Casteel C.L. (2017) A viral protease relocalizes in the presence of the vector to promote vector performance. Nature Communications 8, 14493.

Bergès S.E., Vasseur F., Bediée A., Rolland G., Masclef D., Dauzat M., … Vile D. (2020) Natural variation of Arabidopsis thaliana responses to Cauliflower mosaic virus infection upon water deficit. PLOS Pathogens 16, e1008557.

Bergès S.E., Vile D., Vazquez-Rovere C., Blanc S., Yvon M., Bédiée A., … van Munster M. (2018) Interactions Between Drought and Plant Genotype Change Epidemiological Traits of Cauliflower mosaic virus. Frontiers in plant science 9, 703.

Bhaskara G.B., Nguyen T.T. & Verslues P.E. (2012) Unique Drought Resistance Functions of the Highly ABA-Induced Clade A Protein Phosphatase 2Cs. Plant Physiology 160, 379–395.

Bosu P.P. & Wagner M.R. (2007) Effects of Induced Water Stress on Leaf Trichome Density and Foliar Nutrients of Three Elm (Ulmus) Species: Implications for Resistance to the Elm Leaf Beetle. Environmental Entomology 36, 595–601.

Bouton C., King R.C., Chen H., Azhakanandam K., Bieri S., Hammond-Kosack K.E. & Kanyuka K. (2018) Foxtail mosaic virus: A Viral Vector for Protein Expression in Cereals. Plant Physiology 177, 1352–1367.

Callaway A., Giesman-Cookmeyer D., Gillock E.T., Sit T.L. & Lommel S.A. (2001) THE MULTIFUNCTIONAL CAPSID PROTEINS OF PLANT RNA VIRUSES. Annual Review of Phytopathology 39, 419–460.

Carr J.P. (2017) Exploring how viruses enhance plants’ resilience to drought and the limits to this form of viral payback. Plant, Cell & Environment 40, 2906–2908.

Carr J.P., Donnelly R., Tungadi T., Murphy A.M., Jiang S., Bravo-Cazar A., … Gilligan C.A. (2018) Chapter Seven - Viral Manipulation of Plant Stress Responses and Host Interactions With Insects. (eds P. Palukaitis & M.J.B.T.-A. in V.R. Roossinck), pp. 177–197. Academic Press.

Casteel C.L., De Alwis M., Bak A., Dong H., Whitham S.A. & Jander G. (2015) Disruption of ethylene responses by turnip mosaic virus mediates suppression of plant defense against the green peach aphid vector. Plant physiology 169, 209–218.

Casteel C.L. & Falk B.W. (2016) Plant Virus-Vector Interactions: More Than Just for Virus Transmission BT - Current Research Topics in Plant Virology. (eds A. Wang & X. Zhou), pp. 217–240. Springer International Publishing, Cham.

Casteel C.L., Yang C., Nanduri A.C., De Jong H.N., Whitham S.A. & Jander G. (2014) The NIa-Pro protein of Turnip mosaic virus improves growth and reproduction of the aphid vector, Myzus persicae (green peach aphid). The Plant Journal 77, 653–663.

Castiglioni P., Warner D., Bensen R.J., Anstrom D.C., Harrison J., Stoecker M., … Heard J.E. (2008) Bacterial RNA Chaperones Confer Abiotic Stress Tolerance in Plants and Improved Grain Yield in Maize under Water-Limited Conditions. Plant Physiology 147, 446–455.

Cheng D.-J., Xu X.-J., Yan Z.-Y., Tettey C.K., Fang L., Yang G.-L., … Li X.-D. (2021) The chloroplast ribosomal protein large subunit 1 interacts with viral polymerase and promotes virus infection. Plant Physiology 187, 174–186.

Clough S.J. & Bent A.F. (1998) Floral dip: a simplified method for Agrobacterium-mediated transformation of Arabidopsis thaliana. The Plant Journal 16, 735–743.

Corrales-Gutierrez M., Medina-Puche L., Yu Y., Wang L., Ding X., Luna A.P., … Lozano-Duran R. (2020) The C4 protein from the geminivirus Tomato yellow leaf curl virus confers drought tolerance in Arabidopsis through an ABA-independent mechanism. Plant Biotechnology Journal 18, 1121–1123.

Cui H. & Wang A. (2016) Plum Pox Virus 6K1 Protein Is Required for Viral Replication and Targets the Viral Replication Complex at the Early Stage of Infection. Journal of virology 90, 5119–5131.

Curtis M.D. & Grossniklaus U. (2003) A Gateway Cloning Vector Set for High-Throughput Functional Analysis of Genes in Planta. Plant Physiology 133, 462–469.

Dai A. (2013) Increasing drought under global warming in observations and models. Nature Climate Change 3, 52–58.

Davis T.S., Bosque-Pérez N.A., Foote N.E., Magney T. & Eigenbrode S.D. (2015) Environmentally dependent host–pathogen and vector–pathogen interactions in the Barley yellow dwarf virus pathosystem. Journal of Applied Ecology 52, 1392–1401.

Deng P., Wu Z. & Wang A. (2015) The multifunctional protein CI of potyviruses plays interlinked and distinct roles in viral genome replication and intercellular movement. Virology Journal 12, 141.

Galmés J., Flexas J., Savé R. & Medrano H. (2007) Water relations and stomatal characteristics of Mediterranean plants with different growth forms and leaf habits: responses to water stress and recovery. Plant and Soil 290, 139–155.

González R., Butković A., Escaray F.J., Martínez-Latorre J., Melero Í., Pérez-Parets E., … Elena S.F. (2021) Plant virus evolution under strong drought conditions results in a transition from parasitism to mutualism. Proceedings of the National Academy of Sciences 118, e2020990118.

Guddimalli R., Somanaboina A.K., Palle S.R., Edupuganti S., Kummari D., Palakolanu S.R., … Kavi Kishor P.B. (2021) Overexpression of RNA-binding bacterial chaperones in rice leads to stay-green phenotype, improved yield and tolerance to salt and drought stresses. Physiologia Plantarum 173, 1351–1368.

Guo J., Xu W., Yu X., Shen H., Li H., Cheng D., … Song J. (2016) Cuticular Wax Accumulation Is Associated with Drought Tolerance in Wheat Near-Isogenic Lines. Frontiers in Plant Science 7.

Hikaru S., Hironori T., Fuminori T., Takamasa S., Satoshi I., Nobutaka M., … Kazuo S. (2018) Arabidopsis thaliana NGATHA1 transcription factor induces ABA biosynthesis by activating NCED3 gene during dehydration stress. Proceedings of the National Academy of Sciences 115, E11178–E11187.

Hily J.-M., Poulicard N., Mora M.-Á., Pagán I. & García-Arenal F. (2016) Environment and host genotype determine the outcome of a plant–virus interaction: from antagonism to mutualism. New Phytologist 209, 812–822.

Iuchi S., Kobayashi M., Taji T., Naramoto M., Seki M., Kato T., … Shinozaki K. (2001) Regulation of drought tolerance by gene manipulation of 9-cis-epoxycarotenoid dioxygenase, a key enzyme in abscisic acid biosynthesis in Arabidopsis. The Plant Journal 27, 325–333.

Jež-Krebelj A., Rupnik-Cigoj M., Stele M., Chersicola M., Pompe-Novak M. & Sivilotti P. (2022) The Physiological Impact of GFLV Virus Infection on Grapevine Water Status: First Observations. Plants 11.

Jia H., Zhang S., Ruan M., Wang Y. & Wang C. (2012) Analysis and application of RD29 genes in abiotic stress response. Acta Physiologiae Plantarum 34, 1239–1250.

Kim H., Lee K., Hwang H., Bhatnagar N., Kim D.-Y., Yoon I.S., … Kim B.-G. (2014) Overexpression of PYL5 in rice enhances drought tolerance, inhibits growth, and modulates gene expression. Journal of Experimental Botany 65, 453–464.

Kim W., Iizumi T. & Nishimori M. (2019) Global Patterns of Crop Production Losses Associated with Droughts from 1983 to 2009. Journal of Applied Meteorology and Climatology 58, 1233–1244.

Klein S.P., Schneider H.M., Perkins A.C., Brown K.M. & Lynch J.P. (2020) Multiple Integrated Root Phenotypes Are Associated with Improved Drought Tolerance1 [OPEN]. Plant Physiology 183, 1011–1025.

Krishnaswamy S., Verma S., Rahman M.H. & Kav N.N. V(2011) Functional characterization of four APETALA2-family genes (RAP2.6, RAP2.6L, DREB19 and DREB26) in Arabidopsis. Plant Molecular Biology 75, 107–127.

Laliberté J.-F. & Sanfaçon H. (2010) Cellular Remodeling During Plant Virus Infection. Annual Review of Phytopathology 48, 69–91.

Lamaoui M., Jemo M., Datla R. & Bekkaoui F. (2018) Heat and Drought Stresses in Crops and Approaches for Their Mitigation. Frontiers in Chemistry 6.

Lecoeur J., Wery J., Turc O. & Tardieu F. (1995) Expansion of pea leaves subjected to short water deficit: cell number and cell size are sensitive to stress at different periods of leaf development. Journal of Experimental Botany 46, 1093–1101.

Lellis A.D., Kasschau K.D., Whitham S.A. & Carrington J.C. (2002) Loss-of-Susceptibility Mutants of Arabidopsis thaliana Reveal an Essential Role for eIF(iso)4E during Potyvirus Infection. Current Biology 12, 1046–1051.

Lim S.D.O.N., Lee C. & Jang C.S. (2014) The rice RING E3 ligase, OsCTR1, inhibits trafficking to the chloroplasts of OsCP12 and OsRP1, and its overexpression confers drought tolerance in Arabidopsis. Plant, Cell & Environment 37, 1097–1113.

Liu N., Xie K., Jia Q., Zhao J., Chen T., Li H., … Liu Y. (2016) Foxtail Mosaic Virus-Induced Gene Silencing in Monocot Plants. Plant Physiology 171, 1801–1807.

Livak K.J. & Schmittgen T.D. (2001) Analysis of Relative Gene Expression Data Using Real-Time Quantitative PCR and the 2–ΔΔCT Method. Methods 25, 402–408.

Manacorda C.A., Gudesblat G., Sutka M., Alemano S., Peluso F., Oricchio P., … Asurmendi S. (2021) TuMV triggers stomatal closure but reduces drought tolerance in Arabidopsis. Plant, Cell & Environment 44, 1399–1416.

Martignago D., Rico-Medina A., Blasco-Escámez D., Fontanet-Manzaneque J.B. & Caño-Delgado A.I. (2020) Drought Resistance by Engineering Plant Tissue-Specific Responses. Frontiers in Plant Science 10.

Mei Y. & Whitham S.A. (2018) Virus-Induced Gene Silencing in Maize with a Foxtail mosaic virus Vector BT - Maize: Methods and Protocols. (ed L.M. Lagrimini), pp. 129–139. Springer New York, New York, NY.

Munemasa S., Hauser F., Park J., Waadt R., Brandt B. & Schroeder J.I. (2015) Mechanisms of abscisic acid-mediated control of stomatal aperture. Current Opinion in Plant Biology 28, 154–162.

Murashige T. & Skoog F. (1962) A Revised Medium for Rapid Growth and Bio Assays with Tobacco Tissue Cultures. Physiologia Plantarum 15, 473–497.

Murchie E.H. & Lawson T. (2013) Chlorophyll fluorescence analysis: a guide to good practice and understanding some new applications. Journal of Experimental Botany 64, 3983–3998.

Nawaz G., Lee K., Park S.J., Kim Y.-O. & Kang H. (2018) A chloroplast-targeted cabbage DEAD-box RNA helicase BrRH22 confers abiotic stress tolerance to transgenic Arabidopsis plants by affecting translation of chloroplast transcripts. Plant Physiology and Biochemistry 127, 336–342.

Nemali K.S., Bonin C., Dohleman F.G., Stephens M., Reeves W.R., Nelson D.E., … Lawson M. (2015) Physiological responses related to increased grain yield under drought in the first biotechnology-derived drought-tolerant maize. Plant, Cell & Environment 38, 1866–1880.

Pérez-Bueno M.L., Pineda M. & Barón M. (2019) Phenotyping Plant Responses to Biotic Stress by Chlorophyll Fluorescence Imaging. Frontiers in Plant Science 10.

Prasch C.M. & Sonnewald U. (2013) Simultaneous Application of Heat, Drought, and Virus to Arabidopsis Plants Reveals Significant Shifts in Signaling Networks. Plant Physiology 162, 1849–1866.

Ray S. & Casteel C.L. (2022) Effector-mediated plant-virus-vector interactions. The Plant Cell, koac058.

Revers F. & García J.A. (2015) Chapter Three - Molecular Biology of Potyviruses. (eds K. Maramorosch & T.C.B.T. -A. in V.R. Mettenleiter), pp. 101–199. Academic Press.

Ryals J.A., Neuenschwander U.H., Willits M.G., Molina A., Steiner H.-Y. & Hunt M.D. (1996) Systemic acquired resistance. Plant Cell 8, 1809–1819.

Ryan M.D., King A.M.Q. & Thomas G.P. (1991) Cleavage of foot-and-mouth disease virus polyprotein is mediated by residues located within a 19 amino acid sequence. Journal of General Virology 72, 2727–2732.

Shteinberg M., Mishra R., Anfoka G., Altaleb M., Brotman Y., Moshelion M., … Czosnek H. (2021) Tomato Yellow Leaf Curl Virus (TYLCV) Promotes Plant Tolerance to Drought. Cells 10.

Soma F., Takahashi F., Yamaguchi-Shinozaki K. & Shinozaki K. (2021) Cellular Phosphorylation Signaling and Gene Expression in Drought Stress Responses: ABA-Dependent and ABA-Independent Regulatory Systems. Plants (Basel, Switzerland) 10, 756.

Taiyun W. & Aiming W. (2008) Biogenesis of Cytoplasmic Membranous Vesicles for Plant Potyvirus Replication Occurs at Endoplasmic Reticulum Exit Sites in a COPI-and COPII-Dependent Manner. Journal of Virology 82, 12252–12264.

Taiyun W., Tyng-Shyan H., Jamie M., Jean-François L., Jian H., S. N.R. & Aiming W. (2010) Sequential Recruitment of the Endoplasmic Reticulum and Chloroplasts for Plant Potyvirus Replication. Journal of Virology 84, 799–809.

Team R.C. (2013) R: A language and environment for statistical computing.

Valli A.A., Gallo A., Rodamilans B., López-Moya J.J. & García J.A. (2018) The HCPro from the Potyviridae family: an enviable multitasking Helper Component that every virus would like to have. Molecular Plant Pathology 19, 744–763.

Westwood J.H., Mccann L., Naish M., Dixon H., Murphy A.M., Stancombe M.A., … Carr J.P. (2013) A viral RNA silencing suppressor interferes with abscisic acid-mediated signalling and induces drought tolerance in Arabidopsis thaliana. Molecular Plant Pathology 14, 158–170.

Wilkinson S. & Davies W.J. (2010) Drought, ozone, ABA and ethylene: new insights from cell to plant to community. Plant, Cell & Environment 33, 510–525.

Xu P., Chen F., Mannas J.P., Feldman T., Sumner L.W. & Roossinck M.J. (2008) Virus infection improves drought tolerance. New Phytologist 180, 911–921.

Yamaguchi-Shinozaki K., Koizumi M., Urao S. & Shinozaki K. (1992) Molecular Cloning and Characterization of 9 cDNAs for Genes That Are Responsive to Desiccation in Arabidopsis thaliana: SequenceAnalysis of One cDNA Clone That Encodes a Putative Transmembrane Channel Protein. Plant and Cell Physiology 33, 217–224.

Yamaguchi-Shinozaki K. & Shinozaki K. (1993) Characterization of the expression of a desiccation-responsive rd29 gene of Arabidopsis thaliana and analysis of its promoter in transgenic plants. Molecular and General Genetics MGG 236, 331–340.

Yang X., Lu M., Wang Y., Wang Y., Liu Z. & Chen S. (2021) Response Mechanism of Plants to Drought Stress. Horticulturae 7.

Yu T.-F., Xu Z.-S., Guo J.-K., Wang Y.-X., Abernathy B., Fu J.-D., … Ma Y.-Z. (2017) Improved drought tolerance in wheat plants overexpressing a synthetic bacterial cold shock protein gene SeCspA. Scientific Reports 7, 44050.

Zhou Y., He R., Guo Y., Liu K., Huang G., Peng C., … Duan L. (2019) A novel ABA functional analogue B2 enhances drought tolerance in wheat. Scientific Reports 9, 2887.

